# Optimizing the efficiency of assisted reproduction procedures in the common marmoset (*Callithrix jacchus*) improved IVF outcomes

**DOI:** 10.64898/2026.10.30.763070

**Authors:** Rebecca A. Holton, Joshua T. Brennan, Sophia I. Lipari, James Pickel, Nia Denney, Bhuchitra Singh, Briana Winer-Centurelli, Jinwoo Jun, Yang Zhang, Pan P. Li, Colin D. Robertson, Ingie Hong, Richard Huganir, Reza Kalhor, Christopher A. Ross, James H. Segars, Xiaoqin Wang

**Author notes:** Authors’ Roles XW, CAR, RLH, RK, and JHS acquired funding. RAH, JTB, JP, BWC, BS, JHS, and IH conceptualized and designed the studies. RAH, JTB, SIL, JP, ND, BWC, JJ, CDR, PPL, and JHS developed and performed experimental work and data analysis. All authors provided experimental and methodology feedback. RAH, JTB, SIL, JP, ND, BWC, BS, and JHS performed animal procedures. ND, YZ, and XW managed the marmoset population. XW, JHS, RLH, RK, and CAR supervised the study and design. RAH, JTB, SIL, and JHS wrote the manuscript. All authors contributed to the original manuscript or with editing and reviewing, and approve of the final form. Licensing: CC-BY. The authors have no competing interests to disclose.

## Abstract

The marmoset has emerged as a model species for genetic studies of complex neurodevelopmental conditions because of their brain development, behavior, communication, and compact size. IVF has been used to generate wild-type and transgenic marmoset embryos, but inefficiency of IVF in the marmoset compared to other primates poses challenges for genetic manipulation. To address this limitation, we analyzed the association between variables of age, number of cycles, and progesterone levels for IVF and embryology outcomes. Longitudinal data collected over 2 years indicated that the progesterone levels 24 hours after Cloprostenol sodium (Estrumate) administration, the number of days of ovarian stimulation during an IVF cycle, and the number of oocyte retrievals that an animal had undergone all influenced IVF cycle outcomes. Higher serum progesterone levels at cycle start were negatively correlated with the likelihood of fertilized eggs progressing to the cleavage stage of embryo development. Additionally, a shorter duration of ovarian stimulation (ranging 7 to 14 days of medication) coupled with a GnRH antagonist correlated with an increased percentage of fertilized eggs. We observed a trend that a shorter hormonal stimulation cycle length was associated with a higher percentage of embryos reaching the cleavage stage. The optimization of stimulation cycle protocol and embryology has led to a high percentage of implantation after embryo transfer regardless of whether the embryo was fresh or cryopreserved. These findings suggest that shorter IVF cycles and a progesterone level <10 ng/mL at cycle start were correlated with an increased likelihood of a cleavage stage embryo development.

## Introduction

*In vitro* fertilization (IVF) involves a series of procedures designed to obtain an adequate number of oocytes for subsequent fertilization. Oocyte retrieval is a critical part of this process, requiring techniques that maximize collection efficiency while minimizing procedural risks [1]. The utilization of IVF in the common marmoset (*Callithrix jacchus*) has become increasingly important for advancing translational biomedical research and facilitating the generation of genetically modified nonhuman primate models [2-4]. Compared with Old World primate models, marmosets offer several practical and biological advantages, including small body size, rapid sexual maturation, a short gestation period of approximately 145-150 days, and a high reproductive rate that frequently results in twin offspring [5]. Beyond these reproductive advantages, marmosets possess cognitive and social characteristics that more closely resemble humans than rodent models, making them particularly valuable for studying neurodevelopmental disorders and gene function. While several groups have established protocols for marmoset assisted reproductive technology (ART) and demonstrated the ability to derive genetically modified offspring, a knowledge gap remains regarding strategies to improve efficiency in ovarian stimulation, oocyte maturation, fertilization, and embryo culture [6-8].

Improving the efficiency of IVF in the common marmoset requires identification and optimization of factors influencing oocyte yield and reproductive outcomes. Female age may influence ovarian responsiveness and oocyte quality, while repeated IVF cycles may affect subsequent ovarian responses and oocyte recovery. Additionally, progesterone concentrations may provide insight into reproductive status and the timing of ovarian stimulation and oocyte retrieval, potentially influencing embryogenesis and subsequent maturation. By investigating the associations between maternal age, the number of previous IVF cycles, and progesterone concentrations with IVF outcomes, this study aims to identify parameters that maximize oocyte yield and reduce the need for repeated surgical retrieval procedures.

The establishment and maintenance of a common marmoset colony requires substantial financial and logistical resources, including animal housing, veterinary care, specialized equipment, personnel, hormone stimulation materials, and animal-related procedures. Optimizing ovarian cycle stimulation increases the number and quality of oocytes retrieved, in turn bolstering the embryo yield to facilitate transgenic research. In maximizing the efficiency of these cycles, fewer animals and resources are required, thus supporting the core principle of translational animal research. Given the financial and ethical considerations associated with nonhuman primate research, optimizing IVF and oocyte retrieval protocols is essential to maximizing reproductive success while minimizing resource consumption, reducing the need for repeated procedures and promoting the efficient and responsible use of research animals.

Investigating the effects of maternal age, repeated IVF cycles, and progesterone concentrations on reproductive outcomes in nonhuman primates may provide valuable insights into factors influencing ovarian responsiveness and oocyte yield. Identifying these relationships could inform more effective ovarian stimulation and oocyte retrieval strategies, potentially reducing the number of procedures required to achieve successful fertilization and embryonic development in nonhuman primates.

## Methods

All animal procedures were conducted in accordance with protocols approved by the Institutional Animal Care and Use Committee (IACUC) (Protocol PR24M269). An overview of these methods is presented in Figure 1. This figure was made using Microsoft Copilot on July 24, 2026 [9]. The image was created using the starting prompt: “generate a diagram that shows a roadmap of a typical marmoset IVF cycle, starting with blood draws and progesterone elisa assay to determine progesterone levels, then estrumate to reset their ovary cycle, daily fsh to begin superovulation, cetrotide (a GnRH antagonist) to prevent premature ovulation, oocyte retrieval surgery, sperm collection from a male marmoset to inseminate the eggs (IVF) after in vitro maturation (IVM), fertilization check, microinjection at the zygote or 2C stage to introduce Crispr or lentiviral constructs, continued division to 4C, 8C, 16C, morula, and blastocyst stage, and them embryo transfer into a surrogate marmoset.” The authors have verified the accuracy of the final image.

**Figure 1:**
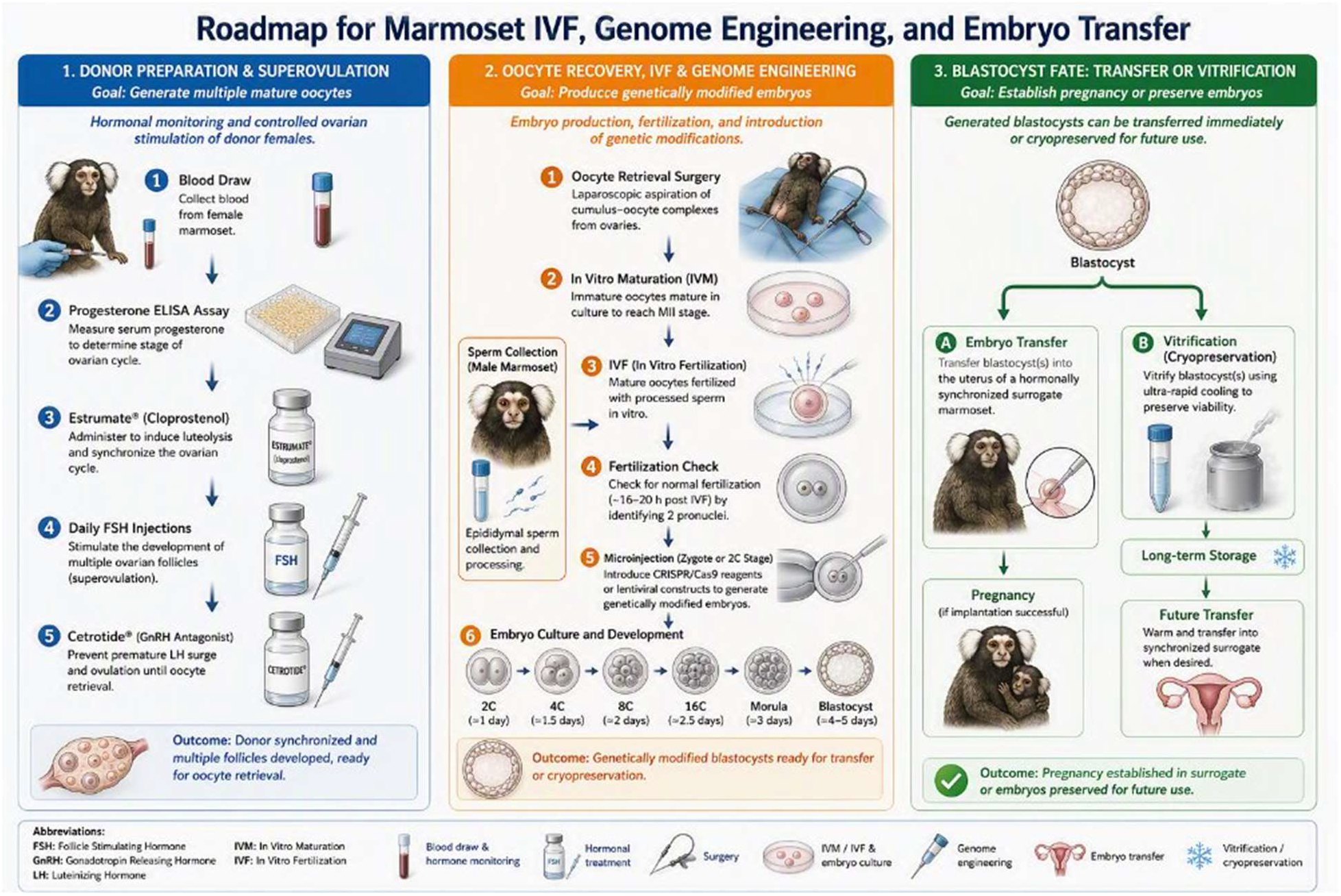
Outline of steps for marmoset IVF and embryo transfer. Roadmap of typical marmoset IVF cycle, including progesterone analysis and hormone administration, oocyte retrieval surgery, sperm collection and IVF, microinjection, and embryo transfer or vitrification. Illustration generated using Microsoft Copilot on July 24, 2026, based on author-provided prompts. The final figure elements and labels were reviewed and verified for accuracy by the authors.

Common marmosets were maintained in single or paired housing under controlled environmental conditions, including a 12-hour light/dark cycle and regulated temperature and humidity. In paired housing, surrogate female marmosets were socially housed with vasectomized males or female offspring to maintain normal social interactions while preventing natural conception. Animals received a standardized Callitrichid Diet 5LK6 (LabDiet), consisting of Vitamin D3 and stabilized Vitamin C and enrichment 3 times per week, including 1 fresh produce, 1 dried food, and 1 protein-rich food with ad libitum access to water. Egg donor ages ranged from 2 to 7 years and surrogate ages ranged from 6 to 13 years.

Following the established protocol from the Sasaki team [4], we measured serum progesterone before and after administering 0.08 ug/mL Estrumate (PGF2α) (MWI Animal Health) to confirm luteolysis before beginning daily ovarian stimulation with 25 IU of 600 IU/mL follicle-stimulating hormone (FSH) (Gonal-F, McKesson). Serum progesterone levels were measured at least twice over the stimulation cycle via Progesterone ELISA Kit (Cayman Chemical Company) and plotted against the provided standards and controls. The marmoset stimulation protocol is depicted in Figure 2, including hormone administration during the follicular phase and ultrasound scans of the uterus and ovaries prior to oocyte retrieval surgery.

**Figure 2:**
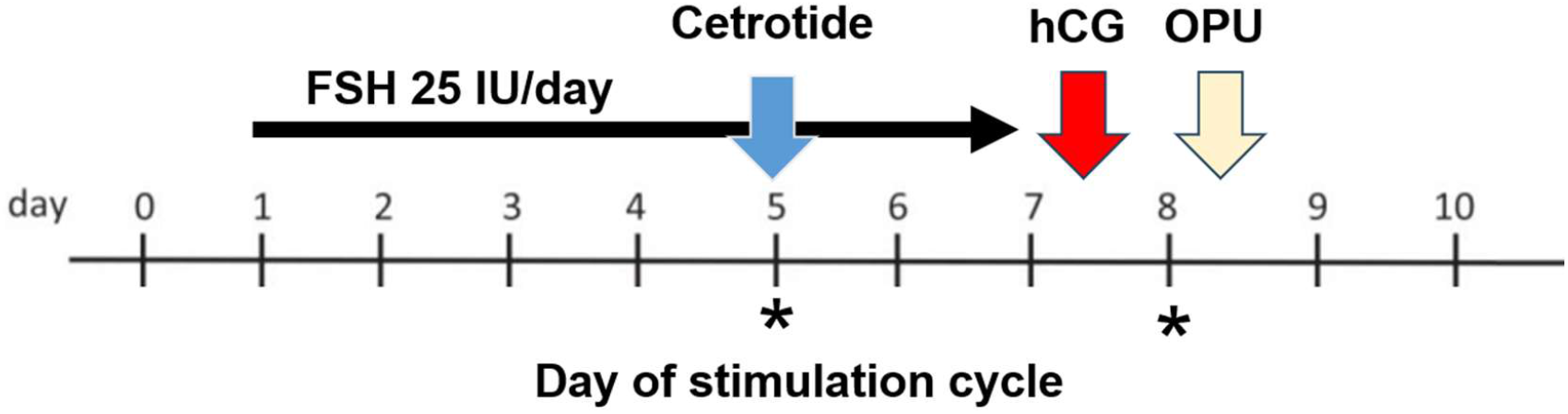
Schematic diagram of an ovarian stimulation for collection of marmoset oocytes. Day 0=Estrumate. Day 1=verification of progesterone level <10 ng/ml. FSH is given at 25 IU/day. Ultrasounds are performed on days as noted by asterisks. hCG is given in the PM on day 7, 18-20 hours prior to the oocyte retrieval (OPU), performed on day 8. Oocytes undergo in vitro maturation before fertilization.

To avoid premature LH surges and ovulation, and to synchronize oocyte maturation, we administered the GnRH antagonist ganirelix acetate (Cetrotide, McKesson) at a subcutaneous dose of 0.125 mg of 0.25 mg/mL during the stimulation cycle, not to exceed 0.1 mg per 100g body weight, followed by 75 IU of 500 IU/mL human chorionic gonadotropin (hCG), administered IM 30-32 hours before egg retrieval (Novarel, McKesson). Follicular development was monitored via ultrasound (Chison Sonobook 9) throughout the stimulation cycle and on the day of retrieval. To ensure surgical candidates had not ovulated prematurely, we evaluated changes in follicle size, the appearance of a corpus luteum, and any increase in post-ovulatory uterine fluid.

Porcine oocyte medium (POM) (CosmoBio, USA) containing heparin was equilibrated overnight and used during the oocyte harvest to aspirate directly from the follicles. During the egg retrieval, oocytes were added to dishes containing additional POM to assess maturation and quality. After sorting appropriately, oocytes were matured overnight in group culture in an Astec IR incubator (38C, 5% CO2, 5% O2) in fresh POM media, supplemented with POM media supplemented with 2% 600 IU/mL FSH, 10% 500 IU/mL hCG and 4% FBS (Corning, 35-010-CV, filter sterilized) to support maturation. The next day, maturation was assessed, and fresh sperm was collected and prepared for IVF in group culture dishes containing Gx-IVF medium (Vitrolife, Fairfax, Virginia). Mature eggs were fertilized with motile sperm (approximately 500 million motile sperm/mL). 18 hours later, fertilization was confirmed and embryos were moved to equilibrated group culture EmbryoGrid dishes (Astec Bio USA) containing cleavage medium (Origio Sequential Cleav, Cooper Surgical) in a CCM iBIS embryoscope (Astec Bio USA) programmed to image each embryo at 12-hour intervals, starting immediately upon placement in the incubator. This provides documented stages of development without disrupting the culturing environment. Approximately 48 hours later, the embryos were moved to equilibrated group culture dishes containing blast medium (Origio Sequential Blast, Cooper Surgical) in a MINC benchtop incubator (Cook Medical) set for hypoxic condition (38C, 5% CO2, 2.5% O2).

Upon successful development, the embryos selected for embryo transfer were brought to the procedure room where a surrogate marmoset was anesthetized and cannulated. Notably, we modified the procedure for embryo transfer. In brief, having experimented with several methods, we use either an aural speculum, or a specially designed mini-vaginal speculum to directly visualize the external cervical os with the aid of a colposcopy at 3x to 15x magnification. Next, the vagina is flushed with embryo culture media. Then, under ultrasound guidance the embryos are transferred with direct visualization aiming for mid-cavity. Custom embryo transfer catheters were provided by Dr. Guoping Feng’s team at Massachusetts Institute of Technology (MIT). The embryos were loaded into a transfer catheter containing 2-3 microliters of blast medium attached to a BD insulin syringe. The catheter was then introduced into the outer cannula until the transfer catheter was located inside the lumen of the uterus, confirmed via concurrent ultrasound and colposcopy. After carefully applying pressure to the plunger, ultrasound and visual inspection of both catheters confirmed delivery into the uterus. Pregnancies were then monitored over the next several weeks, and implantation determined approximately 4 weeks after transfer.

Embryos not selected for fresh embryo transfer were vitrified according to the manufacturer’s instructions (Vit Kit – Freeze NX, Irvine Scientific) or else sequenced to characterize transgenic integration. When needed, the embryos were thawed in accordance with the guidelines included in the warming kit (Vit Kit – Warm NX, Irvine Scientific) and prepared for subsequent embryo transfer or analysis.

### Data Analysis

Statistical analyses and plotting were conducted using GraphPad Prism 11 software. Details of the test used are given in the corresponding figure legends. Data were analyzed using linear regression or Mann Whitney test, where appropriate. A p<0.05 was used to assign significance.

## Results

### Evaluation of Ovarian Stimulation Cycle Factors and their relationship with Embryological Outcomes

Using stimulation cycles defined by prior publications, we collected data from each monkey per stimulation cycle. We then compared conditions to identify parameters that could allow us to increase oocyte yield and decrease egg retrieval surgeries. By analyzing the association of variables of age, number of IVF cycles, and progesterone values with IVF outcomes, we sought to increase the success of marmoset IVF cycles and derive a common, standard protocol for marmoset assisted reproduction.

Blood serum progesterone varies throughout the ovulatory cycle of the marmoset. To correctly time each egg retrieval, treatment of the marmosets with a cloprostenol injection was used to cause luteolysis and reset their cycle to the follicular phase. To proceed with gonadotropin stimulation, blood serum progesterone must decrease post-cloprostenol treatment. We noted that a marmoset was more likely to produce cleavage stage embryos when they had a lower post-treatment progesterone value, particularly when the value was below 10 ng/mL (Figure 3B). Additionally, in subsequent egg retrievals, post-cloprostenol serum progesterone levels increased (Figure 3A), and we noted that animals with fewer surgeries had a more complete luteolysis.

**Figure 3:**
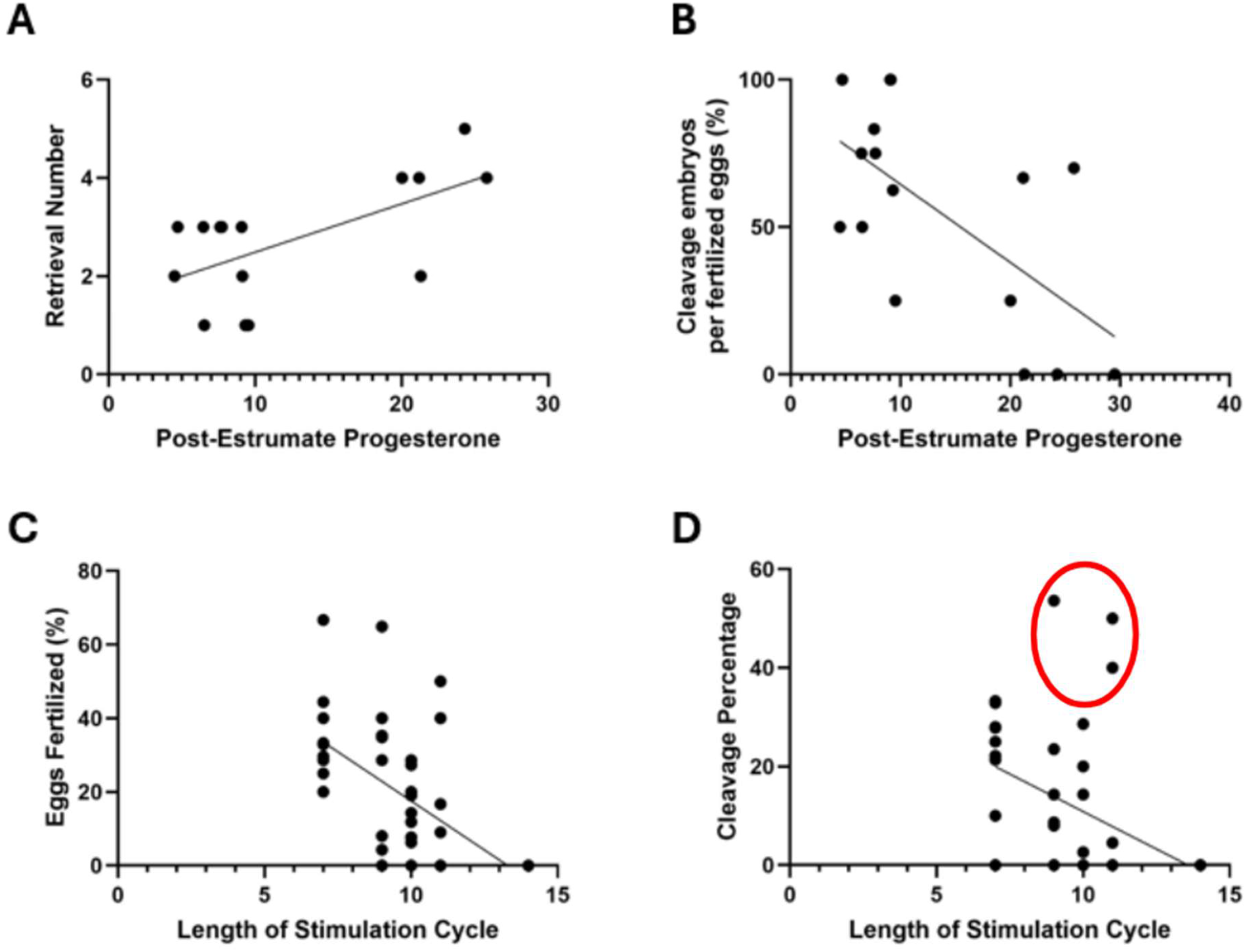
Post-estrumate progesterone and stimulation cycle length were associated with improved embryology in the common marmoset. (A) Progesterone levels at cycle start were positively correlated with the number of retrievals a marmoset undergoes. R^2^ =0.3752, p=0.015, n=15. (B) Increased post-Estrumate progesterone levels reduced the likelihood of embryos developing to cleavage stage. R2=0.4005, p=0.0085, n=16. (C) An increase in hormone stimulation cycle length decreased the percentage of eggs that successfully fertilize via IVF. R2=0.2065, p=0.0025, n=42. (D) An increase in hormone stimulation cycle length may decrease the percentage of cleavage stage embryo formation. R^2^ =0.09122, p=0.051, n=42. Red circle indicates animals treated with GnRH antagonist. Each dot represents one stimulation cycle, n values represent biological replicates. Correlation assessed by linear regression.

We also examined the duration of ovarian stimulation prior to oocyte retrieval. We examined the effect of increasing or decreasing the number of days a female was treated with FSH, length of the stimulation cycle, and assessed the numbers of oocytes retrieved and fertilized to optimize oocyte yield. When compared with the length of stimulation cycle, both the percent of eggs fertilized via IVF and the percent of oocytes that developed into cleavage stage embryos decreased with longer cycle lengths (Figures 3C, D). Notably, there were three instances of unusually high cleavage percentages for their cycle lengths (Figure 3D, red circle). All three of these marmosets were treated with a GnRH antagonist at least 24 hours prior to hCG treatment. The use of a GnRH antagonist was critical for longer stimulation, but shortening the stimulation (Figure 3C) was associated with a high fertilization rate and was cost-saving owing to the reduced number of days animals were treated with FSH. Taken together, these data indicate shorter IVF cycles and a progesterone level below 10 ng/mL at cycle start were correlated with an increased likelihood of a cleavage stage embryo development.

### Changing of Stimulation Cycle Protocol Increased Oocyte and Embryo Yield

Next, we began to restrict hormonal stimulation to animals whose blood serum progesterone levels fell below 10 ng/mL and shortened the stimulation cycle to a consistent eight days (including hCG) and treated all animals with a GnRH antagonist on or about cycle day 5. These changes significantly improved the embryological outcomes. When plotted against subsequent egg retrievals, both the number of good quality oocytes collected per surgery and the percentage of cleavage embryos per fertilized eggs increased (Figure 4A, B). Good quality oocytes were defined as oocytes with an intact zona pellucida that were neither dead nor degenerating at time of collection. In our most recent oocyte retrievals and subsequent embryo culture, we have routinely created embryos that reach compaction and blastocyst (Figure 4C, D), stages that are known to increase pregnancy success when transferred [10]. In our last five oocyte retrievals, we have averaged 24.4 oocytes, with culture that resulted in an average of 7.4 cleavage embryos, and an average of 3.8 of which reach morula stage, and 1.2 that become blastocysts. We observed an improved overall outcome for each stimulation cycle and oocyte retrieval surgery when results were plotted sequentially.

**Figure 4:**
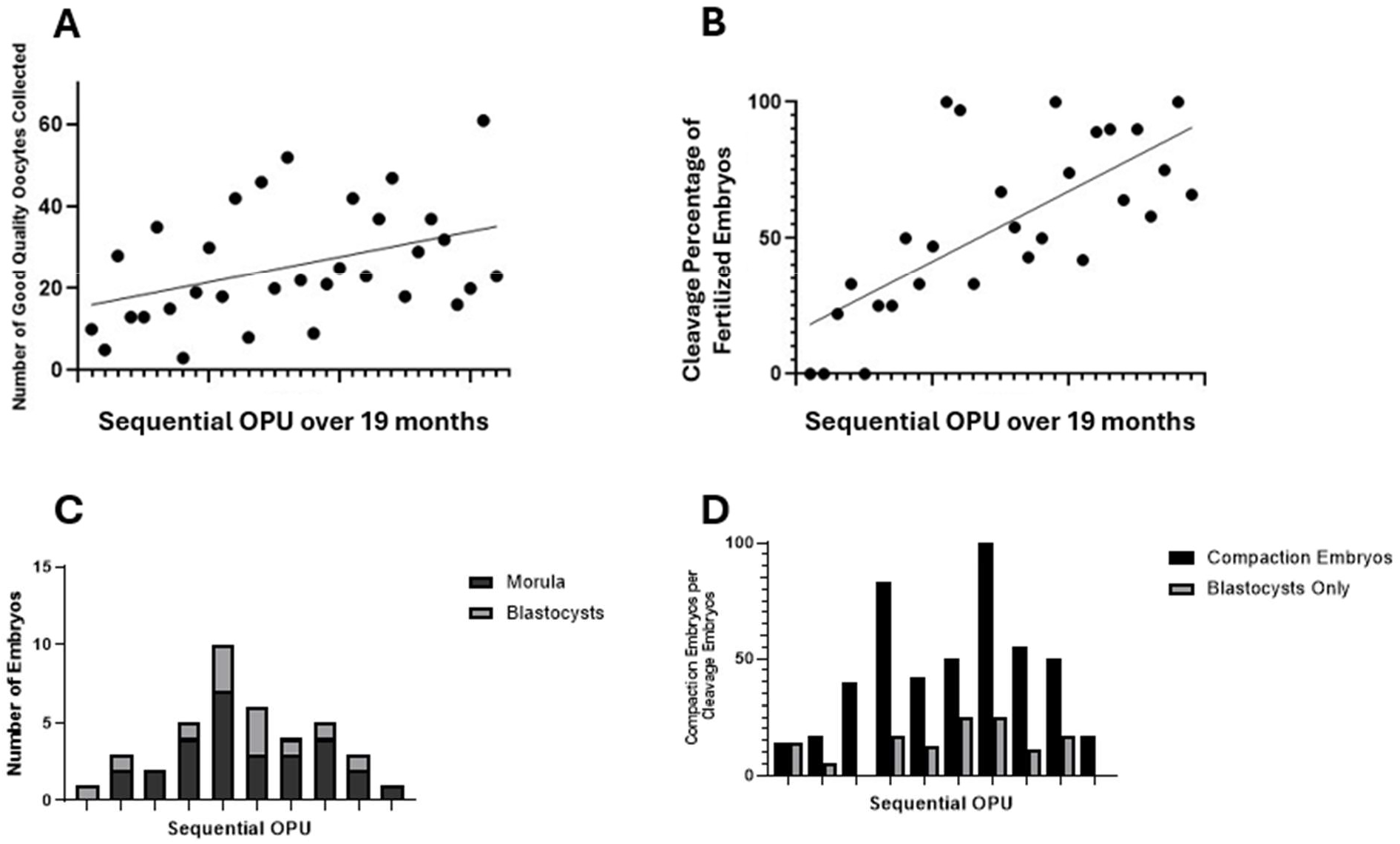
Embryology results plotted versus time. (A) Number of high-quality oocytes increased over time following optimization of cycle stimulation parameters. R^2^ =0.1653, p=0.02, n=32. (B) Embryo cleavage rates improved over time due to changes in stimulation and culture. R^2^ =0.5139, p<0.0001, n=28. Each dot represents one retrieval surgery; n values represent biological replicates. Correlation assessed by linear regression. (C) Number of morulae and blastocysts collected from the most recent 10 oocyte retrievals. (D) Percentage of cleavage embryos that become compaction stage embryos, and the percentage that become blastocysts.

### Embryo Transfer Outcomes

The next step after IVF and embryologic development is the procedure of intrauterine embryo transfer. We typically transferred up to four embryos into parous females, with the transfer day scheduled approximately on or shortly after the animal ovulates. We transferred both fresh embryos and thawed embryos to guarantee the maximum transfer of embryos. Of our fresh transfers, 78% of females displayed signs of implantation during the first eight weeks of gestation. Similarly, 75% of the frozen embryo recipients showed evidence of implantation.

To determine if our embryo vitrification method was satisfactory for marmoset embryos, we compared both implantation rates and lengths of gestation of fresh and cryopreserved groups. Neither parameter showed a significant difference between embryo type. Females with fresh or cryopreserved embryos had a similar likelihood of becoming pregnant (Figure 5A, p>0.99). If a female did show signs of implantation and pregnancy, fresh embryos were likely to survive until week 7.5 of gestation, while thawed embryos survived until approximately gestational week 6. These results were not significantly different (Figure 5B, p=0.74). These data support the belief that our cryopreserved embryos are just as likely to result in a pregnancy as fresh embryos.

**Figure 5:**
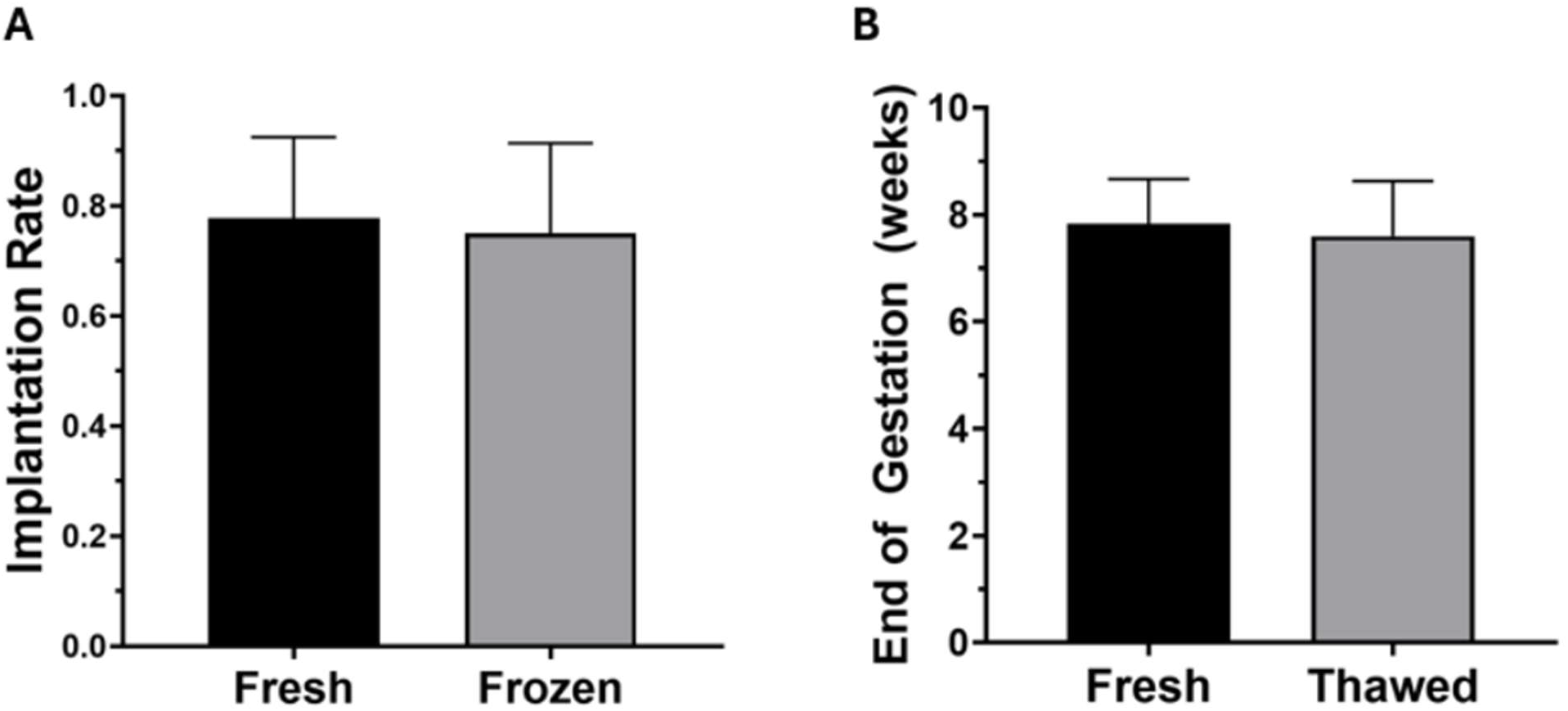
Summary of outcomes of embryo transfers. (A) Implantation rate of transfers with fresh (n=9) and thawed (n=8) embryos. p>0.99. (B) Average length of gestation of transfers using fresh (n=6) and thawed (n=5) embryos. p=0.74. Values represent biological replicates (n=number of transfers). Error bars represent standard error of the mean. Differences compared by Mann Whitney test.

We have successfully carried a wild-type pregnancy to term after an embryo transfer of 4 early-stage embryos into one surrogate female (Figure 6A). We followed the pregnancy using ultrasound imaging to track growth and development of the fetus (Figure 6B). This pregnancy resulted in the birth of a healthy male marmoset (Figure 6C).

**Figure 6:**
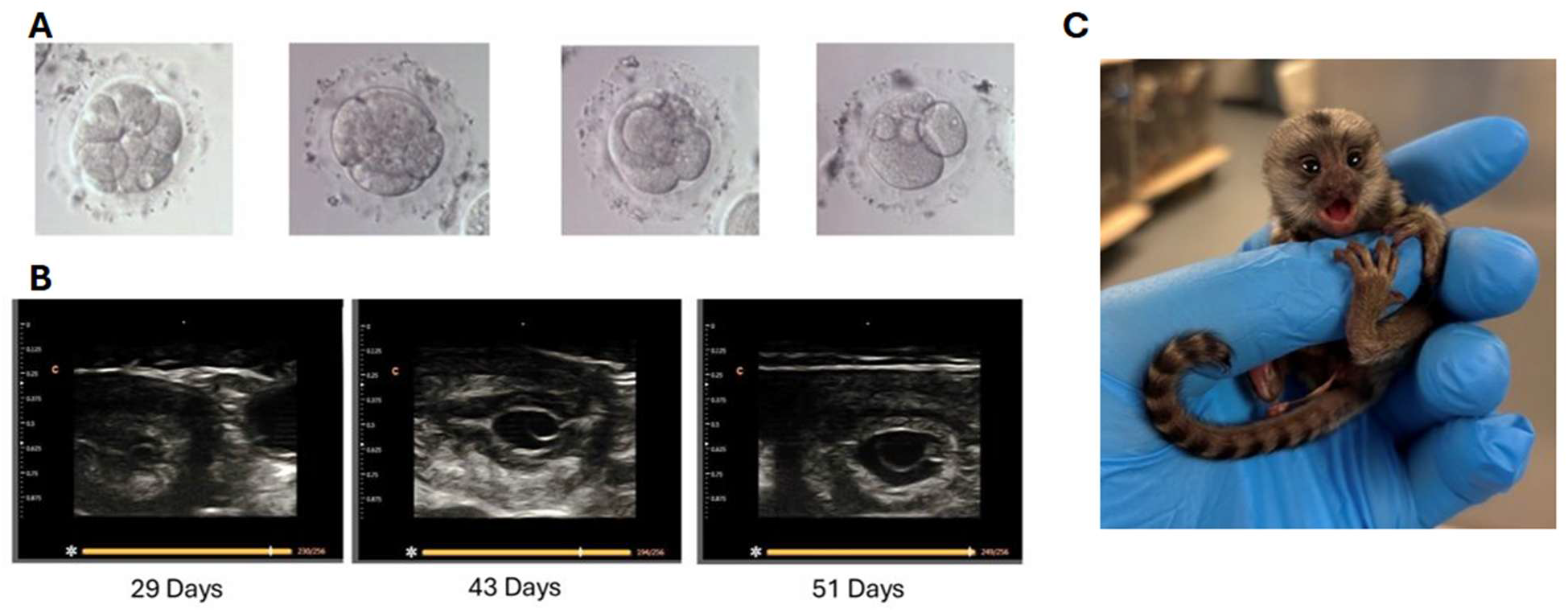
Successful Embryo Transfer and Live Birth. (A) The four embryos transferred into the uterus of a surrogate female. Images captured at 20x magnification. (B) Ultrasound images of the fetus throughout gestation. (C) Resulting neonate offspring of embryo transfer.

We observed that our females benefitted from shorter stimulation cycles after a very low blood serum progesterone level. Additionally, the data suggested that cryopreservation of embryos was a safe and successful method to bank excess embryos for future embryo transfers. Overall, our findings suggest that embryological outcomes can be optimized by tailoring ovarian stimulation cycles to the marmoset colony (Figure 6).

## Discussion

The results indicated that shorter IVF cycles and a post-estrumate progesterone drop below 10 ng/mL were associated with improved embryo development. We also observed trends suggesting a link between treatment with a GnRH antagonist and improved embryo quality. Optimizing each of these steps led to improved ART outcomes regardless of whether the embryo was fresh or cryopreserved. We found that our females produced the best results when their stimulation cycle was between 7 to 9 days, a shift from the duration used by Takahashi and associates [11]. In that study, marmosets were taken to oocyte retrieval on approximately the 11^th^ day of their stimulation cycle, a time point that, in our marmoset population, correlated to a lower fertilization percentage and cleavage percentage unless a GnRH antagonist was used during stimulation. It should be noted that Takahashi et al. showed a similar fertilization percentage to our method, but that study used Intracytoplasmic Sperm Injection (ICSI), rather than traditional IVF. Improving the fertilization rate of traditional IVF enables the use of less invasive, less labor-intensive, and more cost-effective methods for generating successful embryos. The use of traditional IVF also allowed for the natural sperm selection process to occur, thereby avoiding fertilization using sperm with DNA damage or other physiological problems [12].

Commonly administered during the follicular phase in human IVF, gonadotropin-releasing hormone (GnRH) antagonists bind to pituitary GnRH receptors to prevent the luteinizing hormone (LH) surge and resulting ovulation [13]. Premature ovulation during ART is problematic because it causes premature resumption of meiosis, luteinizes the follicles, and reduces oocyte recovery. Thus, premature ovulation leads to unnecessary surgeries for animals undergoing IVF.

An evaluation of the GnRH antagonist, Cetrorelix, observed a 9 to 15-day delay in marmoset ovulation [14]. This delay grants a wider window for retrieving oocytes while preserving egg quality and embryology outcomes. After recovery, Luetjens and associates reported resumption of normal pituitary-ovarian function, indicating that the intended effect causes no long-term damage to the ovarian cycle [14]. By tailoring cycle length and selecting optimal hormone responders before FSH delivery, we minimized reagent use and reduced per-cycle expenses. Additional screening measures allowed us to maximize the individual utility of each animal while also minimizing physical burden and unnecessary stress.

Improving embryology and transfer methods will increase IVF success in marmoset models. Continued optimization facilitates the creation of genetically modified marmosets for disease modeling and streamlines the process for other institutions to customize protocols for their specific colonies. These improvements may, in turn, increase the number of available transgenic marmoset lines and diversify the genes targeted for studies.

## Acknowledgements

We would like to thank the veterinarian team, OR techs, and RAR staff at the Johns Hopkins University School of Medicine for their continued support with our marmoset population. We also thank the Johns Hopkins University School of Medicine Departments of Gynecology and Obstetrics, Biomedical Engineering, Neurobiology, and Neuroscience for their support and assistance with this work.

